# Direct and indirect selection in a proofreading polymerase

**DOI:** 10.1101/2024.10.14.618309

**Authors:** Kabir Husain, Vedant Sachdeva, Riccardo Ravasio, Michele Peruzzo, Wanqiang Liu, Benjamin H. Good, Arvind Murugan

## Abstract

The traits that affect evolvability are subject to indirect selection, as these traits affect the course of evolution over many generations rather than the direct replicative fitness of an individual. However, the evolution of evolvability-determining traits is often difficult to study because putative evolvability alleles often have confounding direct fitness effects of unknown origin and size. Here, we study theoretically and experimentally the evolution of mutation rates in proofreading polymerases with orthogonal control of direct and indirect selection. Mutagenic DNA polymerases enjoy a long-time fitness advantage by enhancing the rate of acquiring beneficial mutations. However, this is offset by a short-time fitness penalty, which we trace to a counterintuitive trade-off between mutation rates and activity in proofreading polymerases. Since these fitness effects act on different timescales, no one number characterizes the fitness of a mutator allele. We find unusual dynamic features in the resulting evolutionary dynamics, such as kinetic exclusion, selection by dynamic environments, and Rock-Paper-Scissors dynamics in the absence of ecology. Our work has implications for the evolution of mutation rates and more broadly, evolution in the context of an anti-correlation between mutation rates and short term fitness.

---

Darwinian evolution is often seen as a tinkering algorithm that incrementally increases the fitness of an organism. As algorithms go, it is remarkably simple: relying on only a few key ingredients — heritable variation and natural selection — to adapt organisms to their environment [1].

However, in contrast to human tinkering, a unique aspect of biological evolution is that the parameters of the algorithm are themselves subject to evolution [2, 3]. The nature and amount of heritable variation available to natural selection is itself under the control of biological factors — ‘modifier’ genes that control the rate at which new variants arise. Examples include genes that affect mutation rates [4–7] and spectra [8, 9], recombination rates [10], horizontal gene transfer, and ‘buffers’ that reduce the impact of deleterious mutations [11–13]. Variation at these loci can influence the rate at which future beneficial alleles arise, thereby altering the dynamics of future evolution.

A particular simple and universally relevant source of variation is the mutation rate, which is set primarily by the biophysics of DNA replication and repair [14]. Genetic changes in the machinery of replication can lead to ‘mutator alleles’: variants with an enhanced mutation rate. Mutator alleles have been found in both laboratory and natural populations. For instance, six of the twelve long-term evolution experiment (LTEE) lineages have fixed a mutator phenotype [7, 15]. In the wild, sampling of bacterial and fungal clades has revealed a large dispersity of mutation rates [16–18]. Even within the restricted timescales of somatic evolution, mutator alleles can have outsize effects, as many cancers are thought to be driven by recurrent genetic instability [19, 20].

Mutator alleles have been the subject of recent theoretical and experimental interest, as a model system to understand when evolution tunes its own parameters. However, determining when and how mutator alleles are favoured by natural selection is challenging. For example, if mutation rates are found to increase in a stressful environment [7, 21, 22], careful work must establish whether this was an inevitable biophysical consequence of the stress, accidental fixation of the mutator alleles, or actually due to the enhanced mutation rate via the acquisition of beneficial mutations. Unfortunately, the evolution of mutation rates is thought to depend on factors that are often difficult to measure in experiments, such as the balance between available deleterious and beneficial mutations [23–32]. As a consequence, it has been difficult to design systematic, well-controlled experiments.

Here, we take advantage of a novel experimental system to study the evolution of mutator alleles. We study the selection of mutagenic alleles of a DNA polymerase in an orthogonal replication system in yeast: the killer DNA virus [33–35]. Here, a DNA polymerase copies only a short cytosolic plasmid contained within the yeast cell. By choosing genes with known fitness landscapes, we control the number and strength of beneficial mutations available under direct selection, and study their consequences on the indirect selection of mutator alleles in the polymerase.

We find that mutator evolution is determined not only by the long-term benefit of mutations that it induces, but also by a short-term cost. This short-term cost — which we trace to a counterintuitive biophysical trade-off between DNA polymerase speed or processivity and mutation rate [36] — grows with the long-term benefit and thus imposes a kinetic barrier to mutator fixation. Even when strongly beneficial mutations are available, mutators can be driven extinct by this short-term cost.

Combining population genetics theory with an engineered, two-gene selection scheme, we map out a phase diagram that predicts when mutators fix. Our results indicate that no one number specifies the ‘fitness’ of a mutator allele when its impact on short-term and long-term fitness are in conflict since they act on different timescales. Accordingly, we find that its population genetics can show ‘rock-paper-scissor’ dynamics. Further, we identify conditions under which ramping up selection strength (e.g., drug levels) over time can select for a mutator, even if no fixed selection strength can do so. We indicate how our results, though established in a concrete experimental system, generalize to evolvability-determining modifier alleles[32, 37, 38] at large, and highlights the fundamentally dynamic nature of fitness when direct and indirect selection are in conflict.

## RESULTS

### A. A minimal experimental system for the evolution of mutator alleles

Prior work on the evolution of mutator alleles has used a small number of mutator alleles that act on the entire genomes of bacteria or yeast [39–41]. Consequently, the number and strength of available beneficial and deleterious mutations — and how they vary in different environments — is unknown and difficult to manipulate experimentally.

We sought a model system in which we could exert rational control over these parameters. We turned to the DNA killer virus found in the yeast species *K. lactis* [33, 42, 43]. This system consists of two cytosolic ‘killer’ plasmids pGKL1 (p1) and pGKL2 (p2), each copied by a dedicated, orthogonal DNA polymerase. Recently, this system was adapted into a directed evolution platform, Orthorep [34, 35], by placing a gene of interest on p1 and linking that gene’s function to host fitness.

Here, we turn this directed evolution platform on its head by using it as an indirect selection platform on the underlying DNA polymerases. We encode a gene with known fitness landscapes (e.g. well-studied antibiotic resistance genes) on p1, select directly on that gene and study the resulting indirect evolution of the DNA polymerase.

Co-opting the OrthoRep directed evolution platform to study the indirect evolution of DNA polymerases provides for several advantages. We can manipulate the strength and availability of beneficial mutations since the p1 plasmid contains a chosen set of genes. The WT DNA polymerase, tp-dnap1, achieves very low mutation rates rates[34] by exploiting kinetic proofreading[44, 45], a broad error-correcting mechanism found across the Central Dogma [46]. As a consequence, mutants of this polymerase can differ in proofreading activity or other aspects of fidelity, resulting in a wide range of mutator alleles, with mutation rates ranging from 10^−7^ substitutions per basepair per generation (sbp) to 10^−4^ sbp [35, 47]. Note that at the mutation rates and timescales studied here, deleterious load is likely to be negligible over the kilobase length-scale of the p1 plasmid.

We first studied the indirect selection of proofreading polymerases during directed evolution of the metabolic gene dihydrofolate reductase (DHFR) from *P. falciparum*. PfDHFR activity is sensitive to the antimarial drug pyrimethamine (pyr), against which it can acquire resistance mutations that have been well-characterized[48, 49]. To carry out this study, we deleted the dihydrofolate reductase (DHFR) gene from the yeast genome and encoded its homolog PfDHFR on p1. The concentration of pyr in our growth media acts as a tuneable selection pressure that acts on a known landscape of beneficial mutations.

We calibrated the system by measuring the strength of a pyr-resistant mutation as a function of pyr concentration. The selection coefficient ranges from *s* = 0.1 without pyrimethamine to *s* = 0.8 at 200 *µ*M pyr, Fig. 1b. Using our experimentally measured data, we simulated a competition between strains with two different mutation rates at different levels of pyrimethamine, at a total population size *N* = 10^7^. Consistent with prior theory [27, 31], the simulation predicts that, at high concentrations of pyr, the mutator allele acquires the beneficial mutation and outcompetes the non-mutator, Fig. 1c.

**FIG. 1.**
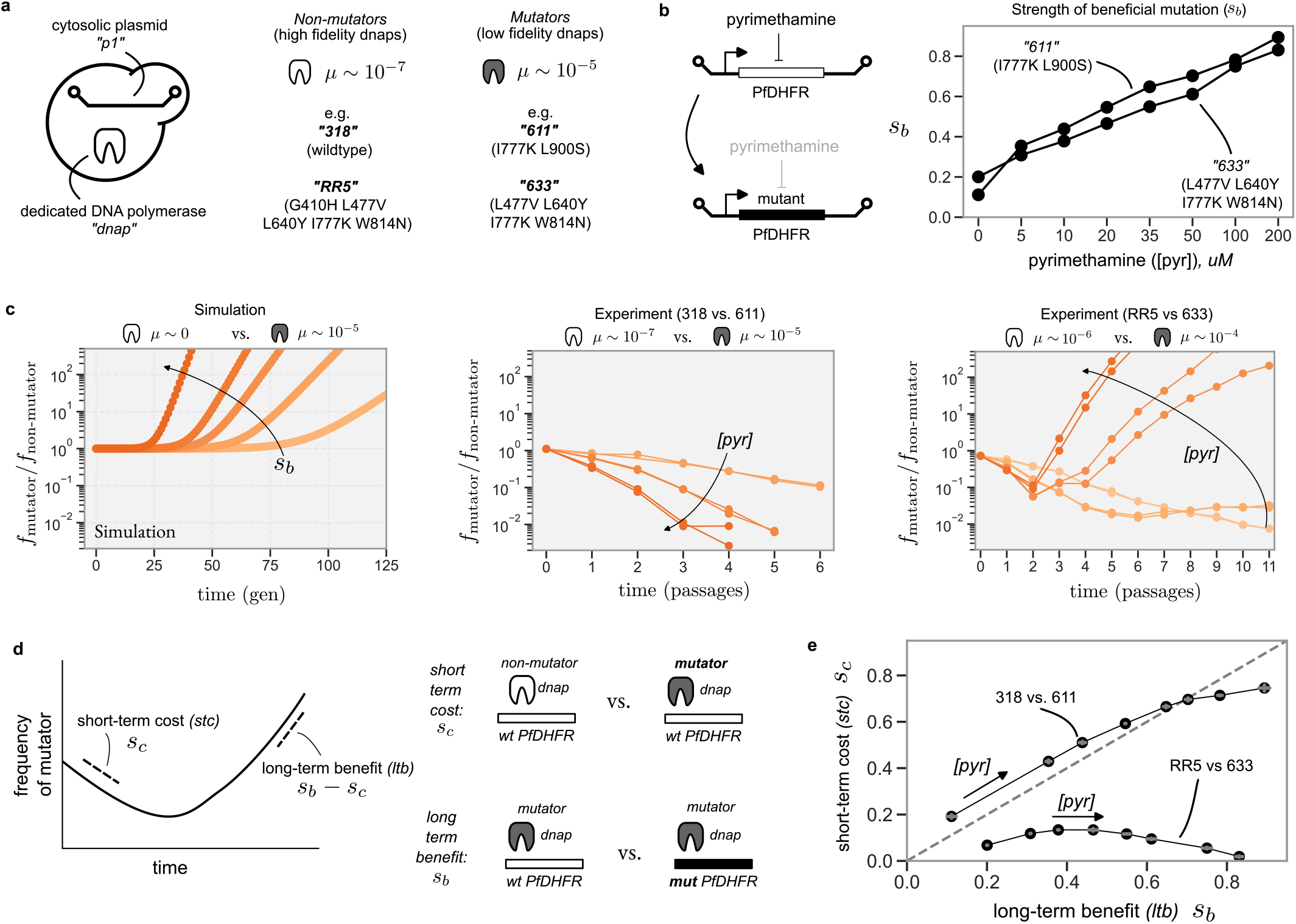
A minimal experimental system for mutator evolution reveals mutators can be driven extinct by a short-time cost that increases with the size of beneficial mutations. **a** In the killer DNA virus that underlies the Orthorep system, a dedicated DNA polymerase (‘dnap’) copies a cytosolic plasmid (‘p1’) independent of the host yeast replication machinery. Indicated are chosen mutator and non-mutator dnaps used in this paper. **b** We encoded the metabolic enzyme dihydrofolate reductase from *P. falciparum* (PfDHFR) onto p1, whose function is inhibited by the drug pyrimethamine. Drug resistant mutations in PfDHFR confer a fitness benefit whose strength varies with pyrimethamine concentration (right). **c** (Left) A simulated competition between a mutator and a non-mutator strain, varying the strength of the beneficial mutation. (Centre + Right) Experimental competition between dnaps 318 and 611 (centre), and 633 and RR5 (right), varying pyrimethamine levels. Two replicates are shown for each pyr level. **d** Definition of the short-term cost (stc) and long-term benefit (ltb). **e** Quantification of stc and ltb as a function of pyrimethamine concentration, for each of the pairs of polymerases competed together in c.

We then performed the same competition in experiment, choosing a pair of polymerases with mutation rates of 10^−7^ and 10^−5^ sbp (termed 318 and 611, respectively; see Fig. 1a). Strains expressing each polymerase were differentially labelled with fluorescent markers mKate and mVenus, mixed 1:1 and passaged daily into fresh media at varying concentrations of pyr. The population composition was monitored by flow cytometry at each dilution.

In contrast to simulation, we observed that the mutator strain 611 was never able to fix in the population. In fact, at higher pyr concentrations, the mutator was out-competed faster, Fig. 1c. We confirmed that beneficial mutations are available to the mutator by competing the pyr-mutant in the mutator 611 background against the non-mutator 318. In this case, the pyr-mutant rapidly fixed in the population at high concentrations of pyr, confirming that selection can act on the beneficial mutation.

We then repeated the experiment with a different pair of polymerases: RR5 and 633. The non-mutator RR5 and mutator 633 had mutation rates of approx. 10^−6^ sbp and 10^−4^ sbp, respectively, similar to the last pair. However, unlike before, we found the mutator does eventually acquire the beneficial mutation and sweep the population at high pyr, Fig. 1c. However, even here the mutator first decreases in frequency before later sweeping the population. The contrast between the two pairs of polymerases, and between the experimental and simulation results, indicated that the mutation rate and the strength of the beneficial mutation are not sufficient to determine the eventual evolutionary outcome.

### B. Short-term costs impose a kinetic barrier to mutator evolution

Why does the mutator 611 lose to 318 even when beneficial mutations are available? We noted that, in all competitions, the mutator population declined at early times. We quantified this ‘short-term cost’ as the early time selective advantage *s*_*c*_ of the non-mutator, Fig. 1d.

In addition, we characterised separately the selective benefit *s*_*b*_ of a beneficial PfDHFR mutation induced by the mutagenic polymerases. Both quantities were found to increase with pyrimethamine, though the short-term cost *s*_*c*_ was markedly stronger for the pair of polymerases in which the mutator eventually lost, Fig. 1e.

We traced the molecular origin of the short-term cost to the processivity of the DNA polymerase. p1 is a multicopy plasmid, with expression of p1-encoded genes increasing with copy number. We hypothesised that our mutagenic polymerases were unable to maintain p1 at a high copy number, and consequently express p1-encoded genes at lower levels with a corresponding effect on fitness, Fig. 2a. Consistent with this, flow-cytometry measurements of a fluorescent protein encoded on p1 showed a sharp reduction in fluorescence when p1 is replicated by the mutagenic polymerases, Fig. 2b.

**FIG. 2.**
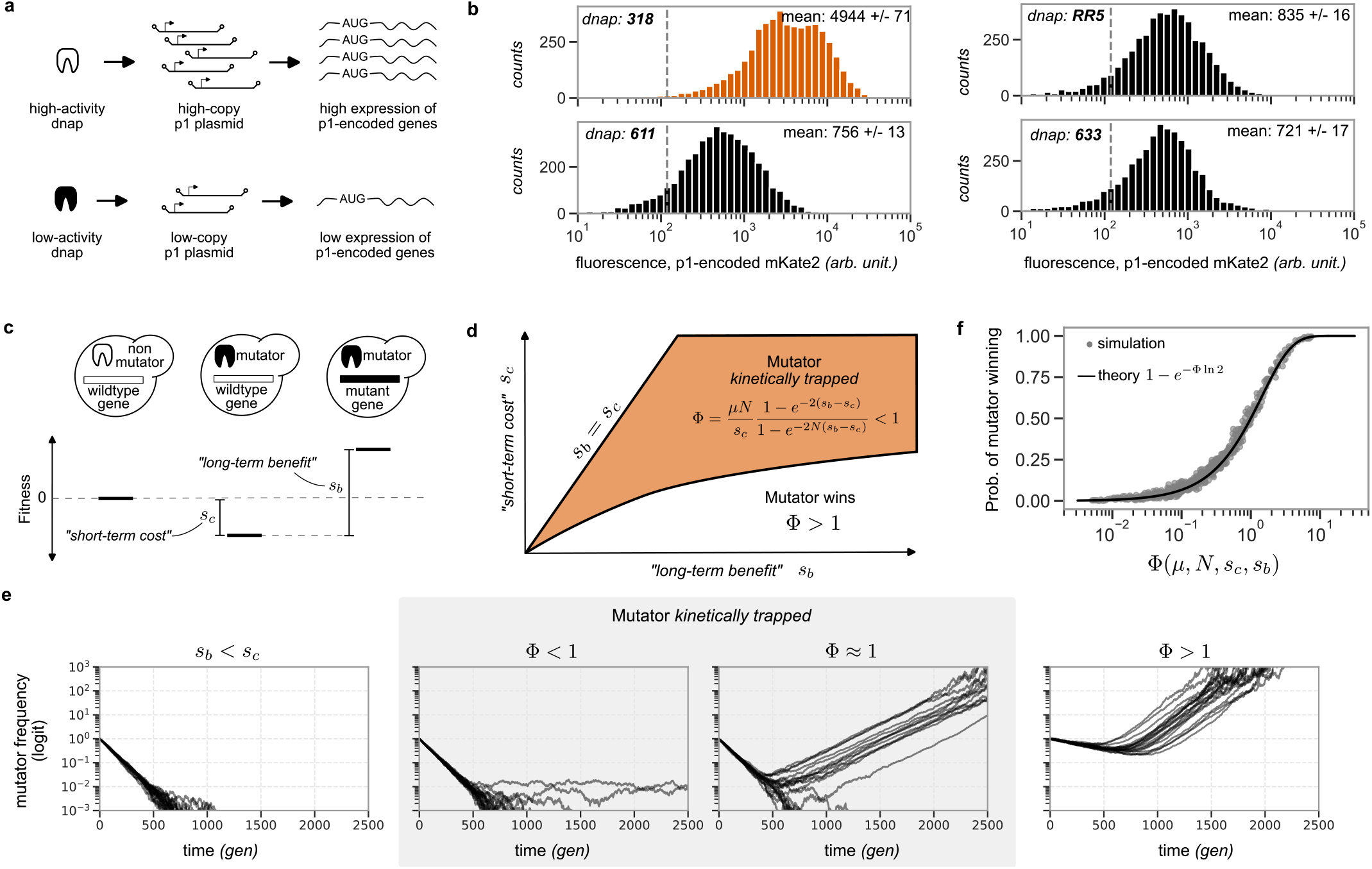
Short-term costs, arising from dnap activity, impose a kinetic barrier to mutator fixation. **a** The activity of the dnap sets the copy number of p1, which determines the expression level of p1-encoded genes. **b** Fluorescence of p1-encoded mKate2 for the four dnaps, measured by flow cytometry. Grey dashed line indicates background fluorescence. Data shows that 318 is a more active dnap than 611, while 633 is comparable with RR5. **c** We calculate the fixation probability of a mutator allele as a function of its short-term cost, *s*_*c*_, as well as the long-term benefit gained from beneficial mutations, *s*_*b*_. **d** A schematic phase diagram, highlighting the kinetic exclusion regime in which mutators lose even though beneficial mutations are sufficiently strong to overcome the short-term-cost. **e** Replicate Wright-Fisher simulations in different regimes of the phase diagram in (d); at Φ ≈ 1, mutator fixation is stochastic, indicating a cross-over rather than a sharp transition. **f** The mutator fixation probability is a universal function of Φ, as shown by a data collapse of many parameter values (grey points) onto a master curve (black).

Differences in expression of PfDHFR also explain why both *s*_*c*_ and *s*_*b*_ grow with pyrimethamine. As pyrimethamine reduces the catalytic rate of the PfDHFR, its effects can be counteracted by a higher expression of the enzyme. Poor expression leads to a higher susceptibility to the drug, and thereby a larger advantage granted by drug-resistance mutations.

Does the short term cost explain the results of our competition experiments? Naively, no. The overall selective advantage of a mutator after it has picked up a beneficial mutation is *s* = *s*_*b*_ − *s*_*c*_. As *s >* 0 for both pairs of polymerases at sufficiently high pyr, we would expect the mutator to have fixed in both cases.

However, this argument neglects the kinetics of the process. As the mutator is being outcompeted by the non-mutator, it only has a finite time ∼ 1*/s*_*c*_ in which to produce a mutant that survives drift and fixes in the population. We refer to this a ‘kinetic barrier’ to mutator fixation, and find that mutators overcome the kinetic barrier when,

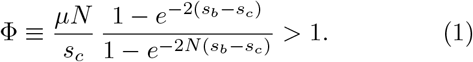

This result is plotted as a phase boundary in the *s*_*c*_–*s*_*b*_ plane in Fig. 2d, and rationalizes the outcomes of our PfDHFR experiments. In one pair of the polymerases (318 vs 611), the high short-term cost paid by the mutator drives the competition into the kinetically excluded regime. Consequently, the mutator is unable to produce a winning mutant before it is driven extinct. In contrast, in the other pair of polymerases (RR5 vs 633), the mutator overcomes this kinetic barrier at high levels of pyrimethamine and is thereby able to fix in the population. (For simplicity, we assumed a zero mutation rate for the non-mutator in this analysis; we relax this assumption later.)

In Fig. 2e we present replicate Wright-Fisher simulations in different regimes of the *s*_*c*_–*s*_*b*_ plane, calculating the probability that the mutator eventually fixes at different values of *s*_*b*_ and *s*_*c*_. Replotting the simulated data as a function of Φ we observe a good data-collapse onto a master curve, confirming the validity of Eq. 1.

### C. Orthogonal control of short term costs and long-term benefits

Thus far, our theory and experimental results make two key predictions. First, that the molecular origin of the short-term cost is the activity of the DNA polymerase. Second, that varying the short-term cost can shift the outcome from the mutator being driven extinct (due to the kinetic barrier), to one in which the mutator is able to acquire beneficial mutations and fix in the population.

These predictions are hard to test in many systems[7, 40] since short and long term effects are hard to disentangle in experiments. In our system, we sought to gain orthogonal control over short-term costs and long-term benefits via a two-gene strategy: one primarily responsible for the short-term cost and the other for the long-term benefit. We integrated the metabolic genes his3 and ura3 on-to p1, with an early stop codon in the open reading frame of ura3 (amino acid 29: caa → taa). The back-ground strain was auxotrophic for histidine and uracil due to disabled his3 and ura3 genes in the genome, and requires expression of these genes from p1 to grow in media without histidine and uracil, respectively (Fig. 3a).

**FIG. 3.**
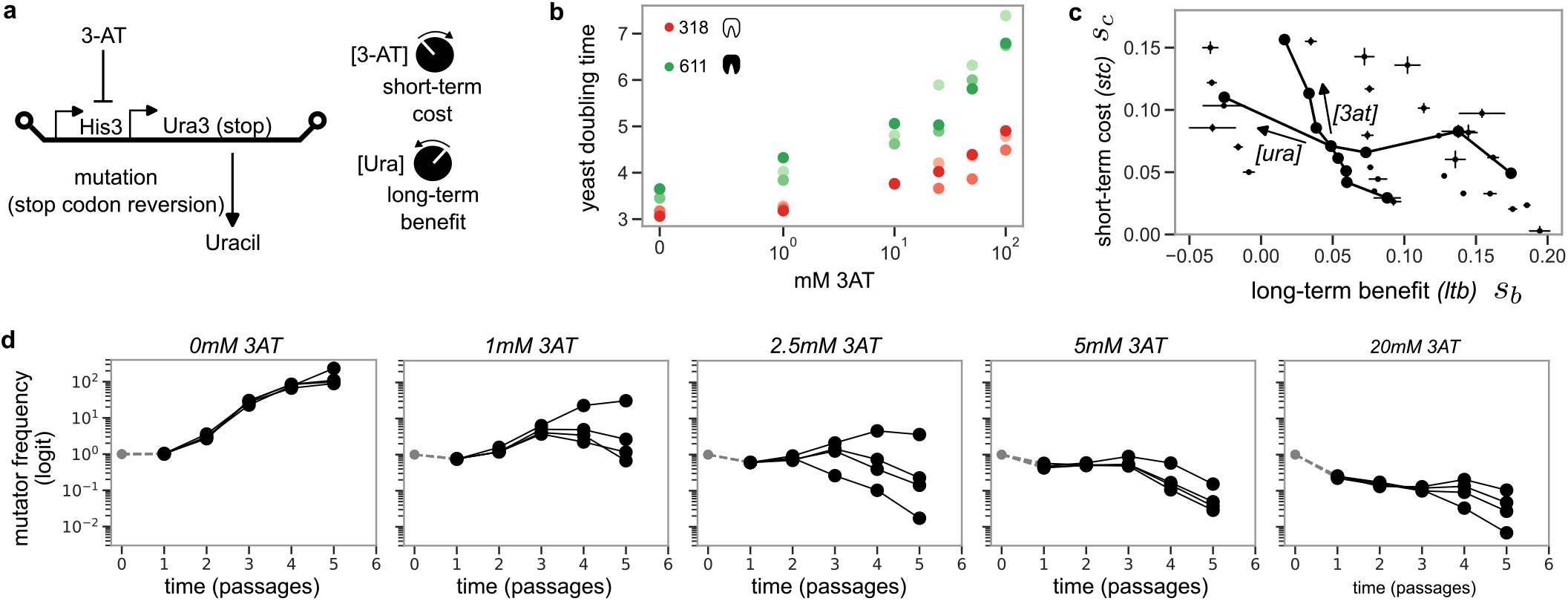
Orthogonal experimental control of short-term costs and long-term benefits reveals different regimes of selection for mutators. **a** Schematic of the his3-ura3* construct on the p1 plasmid, which allows for orthogonal control of short-term costs and long-term benfits by varying 3-AT and uracil levels in the media, respectively. **b** Doubling time of DNA polymerase monocultures in varying levels of 3-AT, showing that the mutagenic, but less active polymerases are more susceptible to the drug. **c** Measured short-term costs and long-term benefits at varying levels of uracil and 3-AT. Solid lines show the effect of varying only uracil or 3-AT. **d** Results from competition between 318 and 611, in depleted uracil media (3.6 g/L) at varying levels of 3-AT.

We reasoned that, in media lacking histidine and supplemented with 3-amino-1,3,4-triazole (3-AT, an inhibitor of his3 function, [50]), the growth-rate of the strain would be a function of his3 expression level, and thereby p1 copy number and polymerase activity. Conversely, removing uracil from the media provides a selective advantage for reversion of the stop codon in ura3. These were borne out in experiment, Fig. 3b. The strain therefore provides orthogonal control over short term costs and long-term benefits by varying the concentration of 3-AT and uracil, respectively.

We confirmed this relationship by measuring the dependence of *s*_*b*_ and *s*_*c*_ on 3-AT and uracil, varying the former between 0 and 50 mM and the latter between 10% and 100% of its usual concentration in our synthetic complete media (i.e. 24mg/L). As expected, and in contrast to our pyrimethamine results, we could now orthogonally tune *s*_*b*_ and *s*_*c*_ by varying uracil or 3-AT concentrations, respectively, Fig. 3c.

With this system in hand, we returned to the polymerases 318 and 611. With PfDHFR as the primary gene, 611 was unable to obtain a beneficial mutation in competition against 318, which we attributed to kinetic exclusion arising from the short-term cost paid by 611. Varying only the concentration of 3-AT, we now observed a transition from a regime in which 611 loses to one in which it picks up a beneficial mutation and fixes in the population, Fig. 3d. We conclude that the observed short term cost is due to differences in the processivity of the polymerase, and that varying the short-term cost is sufficient to alter the eventual fate of the mutator.

Occasionally, at an intermediate value of 3-AT, the mutator’s frequency goes down, up, and finally down again. Such dynamics is likely due to the non-mutator picking up a beneficial mutation at late times and outcompeting the mutator-mutant. Consistent with this, similar dynamics could be observed in simulations with a non-zero mutation rate for the non-mutator. This highlights the complex, non-monotonic population genetics possible with second-order selection.

### D. An inverted speed-accuracy trade-off constrains the evolution of proofreading polymerases

Our results so far result from a conflict between shortterm and long-term fitness effects in mutator evolution. However, this conflict may or may not be relevant to a broad class of mutator alleles. In the particular case of DNA polymerases, 611 and 633 might not be representative mutagenic alleles in that other mutagenic variants might not pay an activity cost. We therefore turned to studying a larger number of DNA polymerase variants, to see if the conflict between short and long-term fitness plays a role more broadly in polymerase evolution.

In Fig. 4 we plot data on 213 polymerase variants from the largest mutational screen of a DNA polymerase accomplished to date, carried out during the development of the OrthoRep platform [35]. Shown is the mutation rate *µ* against the copy number of the p1 plasmid supported by the polymerase — a measure of polymerase activity. Over a broad range, the data shows an inverse correlation between mutation rate and activity: highly mutagenic polymerases are less active, and would be expected to incur a short-term cost against a less-mutagenic, highly-active polymerase such as the wildtype. This result contradicts many prior theoretical expectations [51–55] since proofreading activity[44, 45] is expected to reduce mutation rates at the cost of speed or activity of the polymerase; see [36] for recent work on a potential biophysical explanation for this counterintutive tradeoff.

**FIG. 4.**
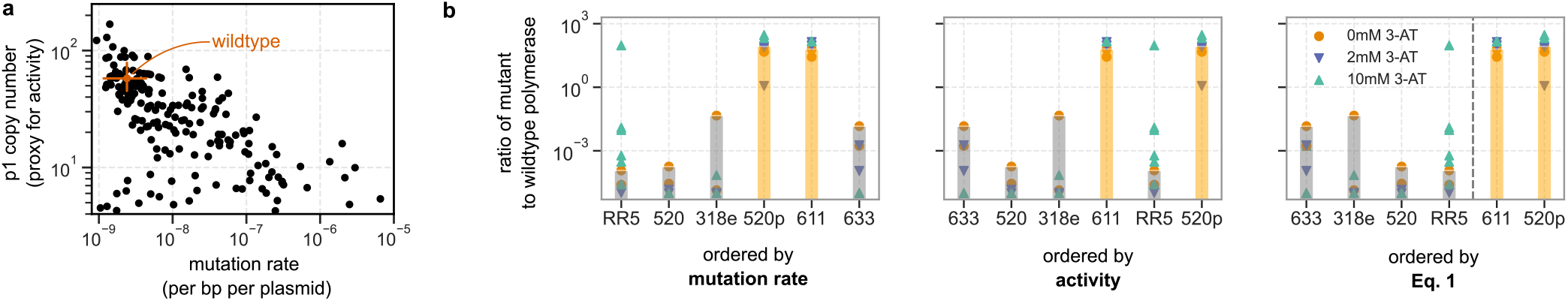
An inverted speed-accuracy trade-off governs the evolution of mutation rates in proofreading DNA polymerases. **(a)** Activity and mutation rate of the killer virus DNA polymerase variants characterized in [35] **(b)** We competed a range of mutant polymerases against the wildtype. Shown is the final fraction of the mutant in the population for each competition, ordered by (i) the mutation rate of the mutant, (ii) the activity of the mutant, and (iii) our criterion, Eq. 1 which correctly rank-orders the competition results.

Setting aside mechanistic origin, here, we focus on the implications for evolution of mutators. The counterintuitive results in Fig. 4 suggest that the evolution of mutagenic polymerases is strongly constrained by short-term costs. If a mutagenic polymerase were to arise in a population, it may be driven extinct despite its higher rate of acquiring beneficial mutations.

To test these ideas, we performed pairwise competition of several polymerase mutants against the wildtype. Six variants were chosen with a range of mutation rates and activities, confirmed by fluctuation assay and p1-encoded fluorescence, respectively. Polymerases were introduced into a strain harboring the his3-ura3* p1 construct and competed against a strain expressing the wildtype polymerase in low-uracil media, thereby selecting for reversion of the stop codon in ura3*.

Fig. 4 shows the frequency of the polymerase mutants in the population after ∼ 40 generations, at which point each mutant is either less than 1% of the population or greater than 99% of the population. We rank order the polymerases by either mutation rate or activity (as assayed by p1 copy number, measured by flow cytometry), to see if either trait can explain the competition results. Neither trait is predictive on its own. While several of the mutagenic polymerases fix against the wildtype, the most mutagenic polymerase is consistently driven extinct. Mutation rate is therefore a poor predictor of mutator fixation. Conversely, activity is also a poor predictor: polymerases 611 and 633-p drive the expression of a p1-encoded gene to similar levels, yet have strongly divergent outcomes in competition against the wildtype polymerase.

In contrast, ordering by the expression Eq. 1 — which takes into account both the polymerase activity and the mutation rate — appropriately separates polymerases that eventually outcompete the wildtype from those that do not – see SI for details. We conclude that the evolutionary fate of mutagenic polymerases depends not only on the long-term benefit of acquiring beneficial mutations, but also on the short-term cost paid in terms of polymerase activity.

### E. ‘Rock-paper-scissors’ dynamics in pairwise competition

Our results so far invoke a tension between short-term costs and long-term benefits in the evolution of mutators. The former acts against the mutator at short times, while the latter favours it at longer times. However, from the perspective of population genetics, can these two factors be combined into a single number that denotes the overall effective ‘fitness’ of a mutator allele?

If such a parameter existed, we could rank-order DNA polymerases by that number into a strict hierarchy (Fig. 5a, left) that predicts outcomes of pairwise competition. In contrast, if instead we can find three mutator alleles whose outcomes in pairwise competition follows a ‘rock-paper-scissors’ dynamic (Fig. 5a, right), no such rank-ordering can exist.

**FIG. 5.**
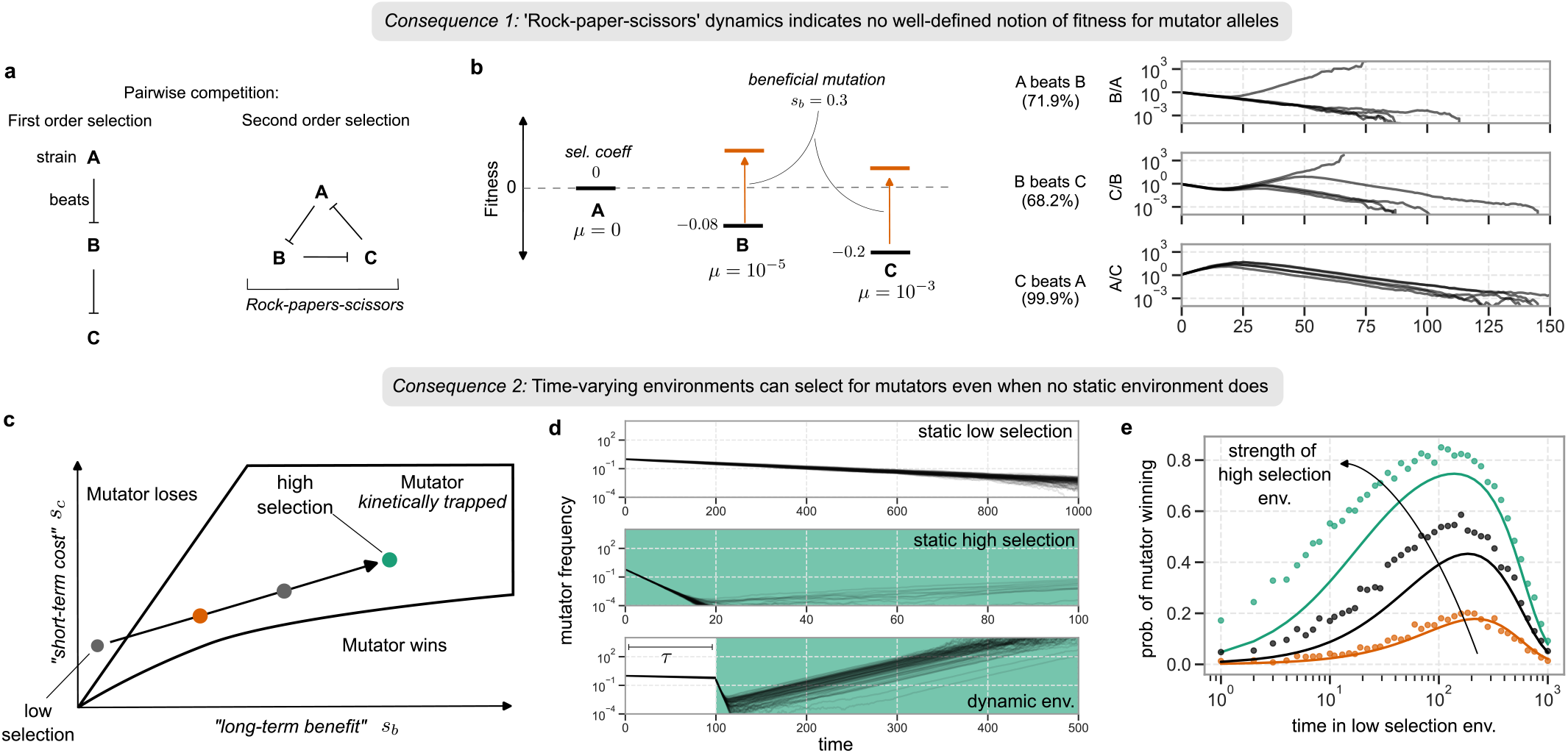
Conflicting short and long term fitnesses lead to unusual evolutionary dynamics of mutator alleles. **a** When genotypes have a well-defined notion of fitness, they can be strictly rank-ordered by pairwise competition outcomes. In contrast, a cyclic ‘rock-paper-scissors’ outcome is inconsistent with any one number characterizing fitness. **b** Simulated pairwise competitions between three strains (Left): a non-mutator (*A* : *µ* = 0, *s*_*c*_ = 0), a medium-mutator (*B* : *µ* = 10^−5^, *s*_*c*_ = −0.08), and a mutator (*C* : *µ* = 10^−3^, *s*_*c*_ = −0.2). *B* and *C* can acquire a beneficial mutation of strength *s*_*b*_ = 0.3. (Right) Pairwise competitions results, with fraction of simulations with indicated outcome written as a percentage. Panels show four representative trajectories from each competition. **c** Schematic showing a permissive, low-selection environment in gray, and three different harsh, high-selection environments in orange, black, and green. All environments are in the regime where Φ *<* 1 (as defined in Eq. 1), in which the mutator loses in a static environment. **d** Schematic of dynamics in static and dynamic environments. In a static permissive environment, the mutator loses as no beneficial mutations are available. In a static harsh environment, mutators are outcompeted due to short-term cost before they can generate beneficial mutations. In a dynamic environment, however, mutators can accrue mutations in the permissive environment that then sweep when they environment becomes harsh. **e** Probability that the mutator fixes in a dynamic environment, as a function of time *τ* spent in the permissive environment. Points show simulated results for three different harsh environment with increasing *s*_*b*_ (as indicated by colour in c). Solid lines are Eq. 2.

In Fig. 5b we present simulations of pairwise competitions between three strains that vary in mutation rate as well as short-term cost (Fig. 5b, left) picked from the scatter plot in Fig. 4a. As we observe in real DNA polymerases, the short-term cost grows with the mutation rate. For each strain, a beneficial mutation of fixed strength is accessible. When mutator *B* is competed against the non-mutator *A, B* is driven extinct by the short-term cost before it can acquire a beneficial mutation. Similarly, though *C* has a higher mutation rate than *B*, both strains are eventually able to acquire the beneficial mutation. As *C* pays a higher short-term cost than *B, C* loses to *B*.

As *B* has lost to *A*, and *C* has lost to *B*, we would expect *C* to lose to *A*. However, we observe that *C*’s mutation rate is so high that it rapidly acquire a winning beneficial mutation and wins against *A*. Thus, *A, B*, and *C* are an example of a ‘rock-papers-scissors’ trio, and the outcome of their pairwise competition cannot be captured by a single ‘fitness’ value. We conclude that no one number summarizes the fitness of a mutator allele.

### F. Mutators are favored by ramping up selection pressure over time

We find that a mutator strain (say strain A) can be selected over a non-mutator (strain B) if the selection pressure on beneficial mutations is ramped up over time, even if strain B is selected over A at any fixed value of selection pressure.

As shown in Fig. 5c, we consider an environments with either fixed low selection or fixed high selection or environments that transition between a low-selection and a high-selection environment. In Wright-Fisher simulations, we find that the mutator loses to the non-mutator if both the fixed low-selection environment and fixed high-selection environment but wins if the environment transitions from low to high selection on a specific timescale *τ* ; see Fig. 5d.

This unusual situation arises from the fact that shortterm cost and long-term benefit have effects that are separated in time. The short-term cost is paid immediately by the mutator, while the long-term benefit is reaped only when (and if) the mutator acquires a beneficial mutation.

As seen in our PfDHFR experiments (Fig. 1), environments in which the long-term benefit of a mutator allele are appreciable are also those in which the short-term costs are high. Consequently, in low selection environments, short-term costs are weak in the low-selection environment, but so are the long-term benefits. In high selection environments, both costs and benefits are high but mutators are driven to extinction by the short term costs before they can accrue a winning beneficial mutation.

By switching environments at the right timescale *τ*, mutators have the opportunity to establish a standing genetic variation of mutants in the low selection environment; these mutants can then be strongly selected in the changed high selection environment. Within a simplified analytical framework, we compute the probability that a mutator fixes as a function of time *τ* spent in the low-selection environment as

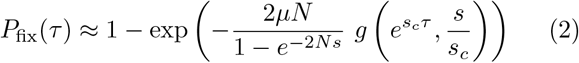

where 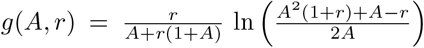, *s*_*c*_ is the short-term cost in the permissive, low-selection environment, and *s* is the fitness of a mutator-mutant in the harsh, high-selection environment.

Eq. 2 is plotted on top of simulation results in Fig. 5d, showing a good agreement and demonstrating that the ‘optimal’ *τ* is ∼ 1*/s*_*c*_. We conclude that the temporal separation between short-term costs and long-term benefits allows the population genetics of the mutator to ‘resonate’ with environmental timescales, highlighting the unusual dynamics of second-order selection.

## DISCUSSION

Our results show that that the evolution of mutation rates in polymerases is determined by a conflict between two traits - mutation rate and replication activity (e.g., speed or processivity). However, unlike other common cases of pleiotropy where one allele affects multiple traits, the two traits here manifest at different timescales. One trait — the mutation rate — is subject only to second-order selection that acts on the lineage, not the individual. The other trait – replication activity — is subject to first order selection, as it determines the replication speed of the individual’s genetic material.

If the two traits had been in concert, the above distinction would not have a qualitative impact. While our case study involved proofreading polymerases, the kind of pleiotropy seen here — negative direct costs associated with positive first order costs — should be expected to be seen just as often as the converse. For example, codon usage can change the availability of beneficial mutations and mutational robustness, a second order effect; but codon usage also typically has first order direct effects in terms of protein translational efficiency. These effects could be linked in conflicting directions.

Similarly, deleterious load, while mechanistically distinct from the direct cost in replication activity studied in this work, could effectively play a similar role as seen here[27, 31, 56]. In particular, if the positive and negative tails of distribution of fitness effects tend to scale together, higher mutation rates will be linked to higher deleterious load and more strongly beneficial mutations, much like in the work here. Consequently, the counter-intuitive evolutionary consequences such as rock-paper-scissors and environmental timescale-dependent selection of mutators might be seen in a range of contexts beyond proofreading polymerases.

Our work suggests the intriguing possibility that mutation rates could ‘ratchet’ or lock in place at unusually low mutation rates. The trade-off in Fig. 4a suggests drift towards higher mutation rate would be selected against due to loss of activity which appears to be significant for the WT polymerase here. As a consequence, drift dynamics would be ratchet-like, with many more non-deleterious mutations available towards lower *µ* than towards higher *µ*. In fact, this possibility could offer an alternative explanation for why the mutation rate of the killer DNA virus exploited here is so low, adding a layer to the drift-barrier hypothesis[3, 28]. The WT mutation rate is *µ* ∼ 10^−9^ substitutions per base while the WT p1 plasmid is of length *L* ∼ 10^4^ bases. The resulting deleterious load *µL* ∼ 10^−5^ ≪ 1 is weak; our quantitative results in this study (e.g., Fig.4) suggest that the need for sufficient replication activity exerts a substantially stronger selective pressure than deleterious load in maintaining such unusually low mutation rates.

## ACKNOWLEDGMENTS

We thank the Kavli Institute for Theoretical Physics, University of California, Santa Barbara (UCSB), where some of this work was done. AM acknowledges support from the NIGMS of the NIH under award number R35GM151211 and the NSF Physics of Living Systems (PHY-2310781). BHG is a Chan Zuckerberg Bio-hub - San Francisco Investigator, and acknowledges support from the NIGMS of the NIH (award number R35GM146949) and the Alfred P. Sloan Foundation (grant FG-2021-15708). This work was supported by the National Science Foundation through the Center for Living Systems (grant no. 2317138).

## Supplementary Information

### Strains, growth conditions, and plasmids

Yeast were grown either in YPD (1% yeast extract, 2% peptone, 2% dextrose) or in synthetic complete (SC) media (1.7 g/L yeast nitrogen base, 5 g/L ammonium sulfate, 36 mg/L myo-inositol, and 2g/L of a homemade amino acid mix) with the appropriate drop-outs. Solid media included 2% agar.

The base OrthoRep strains GA-y103 and GA-y319, as well as plasmids containing the tpdnap1 variants 318, 520, and 633, the PfDHFR gene, and p1 integration cassettes were a kind gift from Chang Liu. Other tpdnap1 variants were constructed by site-directed mutagenesis. To monitor population composition in competition experiments, we cloned a fluorescent protein onto the same plasmid as the polymerase – either mKate or mVenus.

Genomic deletions were performed by CRISPR/Cas9. Guide RNA sequences were cloned into the Cas9 plasmid by blunt end ligation, while 100bp double-stranded repair templates containing 50bp of homology upstream and downstream of the desired ORF were made by PCR. Each knockout was done with 500ng to 1ug of pCas plasmid and 1 to 5ug of repair template; knockouts were verified by diagnostic PCR.

Unless otherwise noted, all cloning was done with a home-made Gibson assembly mix.

To integrate genes onto p1, the desired ORFs were first cloned into integration cassette plasmids. Linear integration cassettes, containing flanking regions of homology to p1, were amplified by PCR and transformed directly into yeast. Transformants were restreaked to single colonies, and successful integration confirmed by PCR.

### Flow cytometry and quantification

Flow cytometry was done on Attune Nxt or Agilent Penteon instruments, flowing > 10^5^ cells for each measurement. To identify mKate-positive and mVenus-positive cells in competition experiments, thresholds were manually set and verified for all experiments. > 80% of all cells were positive for either mKate or mVenus; a small fraction of flowed events were positive for both or neither – these were discarded from the analysis.

To quantify the copy number of p1, we transformed polymerase variants into a strain with a p1-encoded mKate. Cells were passaged 1:100 twice to stabilise p1 copy number, then back-diluted and grown to log phase for measurements. A non-fluorescent yeast strain (of the same genetic background) was flowed alongside as a negative control.

### Fluctuation assays

The mutation rate of all polymerases used were measured by Luria-Delbrück fluctuation assays, using the reversion of a stop codon in either p1-encoded leu2 or ura3 as a selectable marker. Assays were set up with ∼ 20 replicate cultures each, and mutation rates were inferred by maximum likelihood fitting of the Luria-Delbrück distribution.

### Competition time series

Pairwise competitions were conducted in deep-well (96-well) plates, with populations propagated by passaging into fresh media and composition monitored by flow cytometery at every passage. Competitions were initiated by growing monocultures of the mutator and non-mutator strains (i.e. 611 vs 318 for both PfDHFR and HisUra, 633 vs RR5 for PfDHFR alone) to saturation, and mixing 1:1 as assessed by OD600. The mix of cells was then diluted (10uL into 400uL for PfDHFR strains, 1 uL into 1mL for HisUra strains) into wells of a deep-well plate, into media containing the appropriate selective media (i.e. dropout media supplemented with varying amounts of pyrimethamine for PfDHFR, or with 3-AT and uracil for HisUra strains). Each selective condition was run with at least two independent replicates in different wells of the plate.

Competitions were then grown without shaking for approximately 2 days, at which point they were passaged identically into the same selective media. A portion (∼ 100 uL) of saturated culture was removed for analysis by flow cytometry at every passage. Competitions were terminated when one strain exceeded the other by over a hundred-fold.

**Figure 1:**
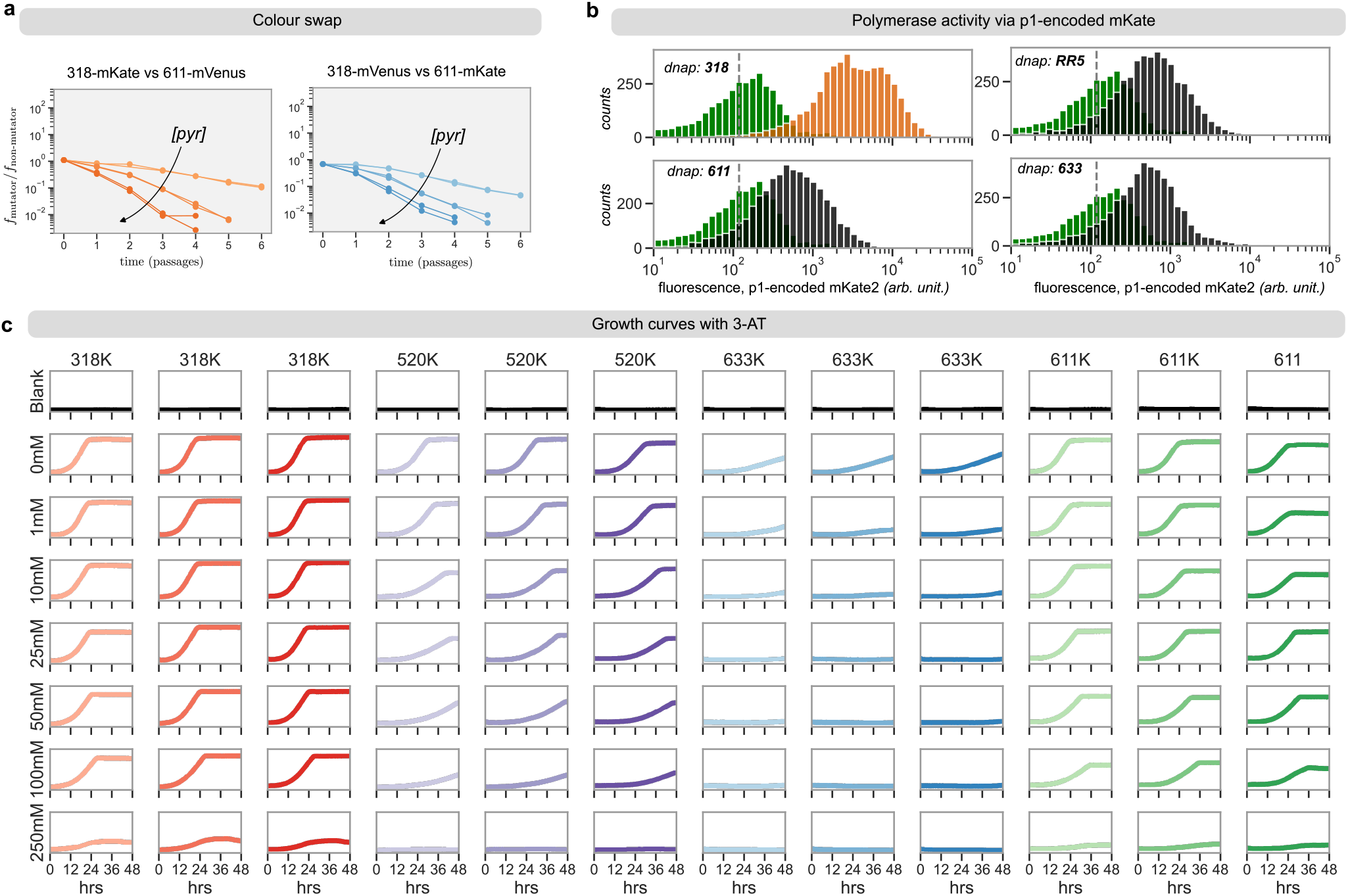
**a** Colour swap competition experiment with 318 and 611 in the PfDHFR strain. **b** Fluorescence of p1-encoded mKate2 for the four dnaps, measured by flow cytometry. Grey dashed line indicates mean background fluorescence, green is the distribution of background fluorescence. **c** Growth curves of indicated dnap at varying 3-AT levels.

To ensure that the choice of fluorescent protein does not affect our conclusions, we repeated our 318-611 PfDHFR competitions with ‘colour-swapped’ strains, Fig. 1. We observe qualitatively similar dynamics.

### Measuring short-term-costs and long-term-benefits

We measured short-term-costs and long-term benefits by pairwise competition, using the mKate- and mVenus-marked strains described above. Briefly, the chosen strains (see below) were normalised by OD600, combined 1:1, and analysed by flow cytometry to obtain the initial ratio of strains. The mix of cells was then diluted either 1:40 into 400uL of the appropriate dropout media supplemented with pyrimethamine (PfDHFR strains), or 1:1000 into 1mL SC-HWU supplemented with 3-AT and uracil (His-Ura strains) – in wells of a 96-well plate. Each growth condition was performed in at least two replicates. Cells were then grown at 30C for approximately two days, following which each competion was analysed by flow cytometry to obtain the final ratio of strains.

To measure short-term costs, we competed together a mutator with a non-mutator (i.e. 611 vs 318 for both PfDHFR and HisUra, 633 vs RR5 for PfDHFR alone). Denoting by *r* _*f*_ (*r*_0_) the ratio of mutator to non-mutator at the final (initial) time, the selection coefficient is

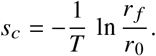

where the number of generations, *T*, was estimated from the dilution factor used to set up the competitions: i.e., 5 for PfDHFR strains and 10 for His-Ura strains.

To measure the long-term-benefit *s*_*b*_, we first obtained mutants in the mutator background by either passaging repeatedly in liquid culture containing pyrimethamine (for PfDHFR) or by selecting on an SC-Ura plate (for the His-Ura strain). These were then competed against the non-mutator as described above. The corresponding selection coefficient, *s* = *s*_*b*_ − *s*_*c*_, was computed as:

**Table 1:**
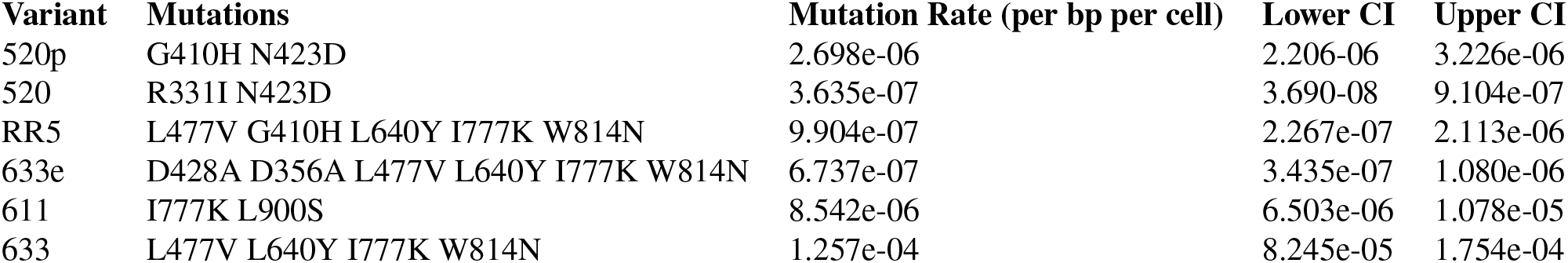
Mutation rates of polymerase variants measured in this study.

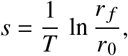

where *r* _*f*_ and *r*_0_ are as before, except now for the mutator-mutant. *s*_*b*_ was then obtained by adding *s*_*c*_ to this.

### Competing mutant polymerases against the wildtype

We competed six mutant polymerases against the wildtype. Competitions were done as above, with the following modifications. Cells were grown in SC-HWU supplemented with 10% uracil and either 0 mM, 2 mM, or 10 mM 3-AT. The competitions were passaged 1:100 in 400uL of media, and the population was analysed by flow cytometry after the first, second, and seventh passage – after which the experiment was terminated. The only exception was RR5 at 10 mM 3-AT, which had to be re-done due to human error and was passaged only up to the fourth passage.

To rank order competition results, we measured mutation rates with fluctuation assays as described above. To approximate the short-term cost, we measured the expression level of a p1-encoded fluorescent protein (mKate) replicated by each of the mutant polymerases; as higher expression level corresponds to a lower short-term cost, we approximated the latter as ∝ 1/expression level.

To rank order by our theory (Eq. 1 in the main text), we further assumed that the fitness of the beneficial mutant (*s*_*b*_ − *s*_*c*_) was the same for all variants (as competitions were done in severe uracil depleted conditions, a ura+ strain would have a strong fitness advantage). Using the simplified criterion (Eq. 4 of this SI), we therefore computed *μN*/*s*_*c*_ using the expression level to approximate *s*_*c*_ as above – giving us the final rank ordering shown in Fig. 4b of the main text.

### Growth curves

In Fig. 3 of the main text, we report the doubling times of different polymerases at varying levels of 3-AT. These were obtained from growth curves in a 96-well plate. Briefly, saturated cultures of HisUra strains with different polymerases were diluted 1:200 in SC-HW supplemented with varying amounts of 3-AT. These were used to seed a clear 96-well plate, with each condition run in triplicate. Twelve wells were filled only with empty media to serve as blanks. Plates were grown at 30C with shaking in a BMG Spectrostar Nano, taking OD600 readings every 20 minutes. Data was background subtracted and fit in the exponential growth phase; doubling times were computed from the fitted growth rate.

### Kinetic trapping and mutator fixation

In the main text, we introduce ‘kinetic trapping’, in which mutators are driven extinct even though beneficial mutations are available. To see this, we compute here the probability that a mutator strain gains a winning mutation and fixes in the population before being driven extinct.

For simplicity, suppose that the non-mutator has a mutation rate of 0, and that the mutator and non-mutator are at equal proportion at the initial time *t* = 0. For the mutator, mutations occur with rate *μN f*_mut_(*t*), where *f*_mut_ is the mutator frequency, and each mutation has a probability

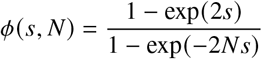

of fixing, with *s* = *s*_*b*_ − *s*_*c*_. Putting it all together,

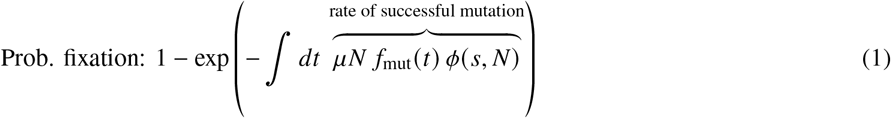

The fraction of mutators in the population satisfies

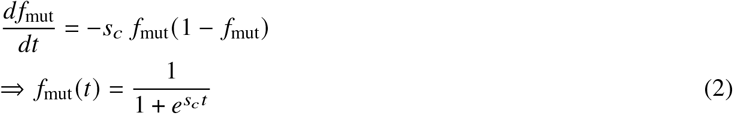

with an initial condition of *f*_mut_(*t* = 0) = 1/2. Plugging everything in, we get:

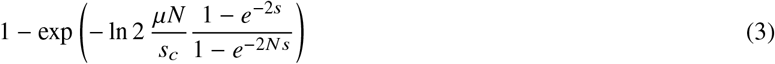

In the main text, we report the quantity in the exponent as Φ. If we expand Φ, up to O(1) factors, we obtain a simpler, approximate criterion for mutators winning:

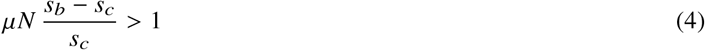

### Time-varying environment (“ramp”)

Let us suppose we go between two environments:

- Env. 1 (low selection): Short-term cost *s*_*c*_ and long-term benefit 0 (ie mutator-mutants have the same fitness as mutator-wildtype).
- Env. 2 (high selection): Very high short term cost (we’ll never need it explicity), but a long-term benefit such that the overall fitness of a mutator-mutant is *s* > 0.

By definition, env. 1 is in the bottom-left region of the phase diagram in Fig. 2 of the main text. For simplicity, we will suppose that env. 2 is in the centre (‘kinetically trapped’) phase. Consequently, mutators do not win in either static env. 1 or in static env. 2 (or any environment that interpolates between them)

The population will start in env. 1 with an initial frequency of mutators *f*_mut_ = 1/2, spend time *τ* there, and then find itself in the harsh env. 2 for the remainder of the calculation. We will assume that the only route to fixation is that a mutator-mutant arises in env. 1, survives until time *τ*, and then fixes in env 2. This neglects the possibility of successful mutants arising in env. 2 itself – as env. 2 is in the kinetically trapped phase, this is a low-probability event, and so we neglect it.

Once again, we will calculate fixation probability from the rate of generating successful mutants:

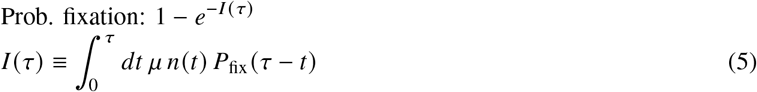

except now the probability that a particular mutation fixes depends on it surviving until the environmental switch. We can write *P*_fix_ (*t*) as follows. First: we’ll need to average over the frequency of the lineage established by the mutant at time *τ* when the environment switches:

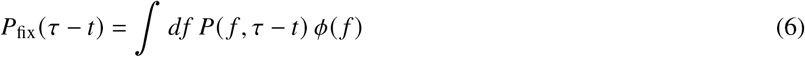

where 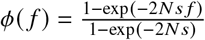 is the prob. that a mutant with frequency *f* fixes, and *P*(*f, τ* − *t*) is the probability that a mutation that occured at time *τ* − *t* in env. 1 has frequency *f* at time *τ*:

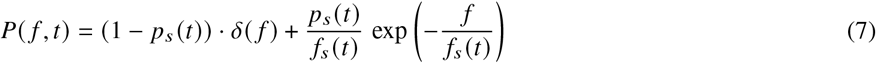

where *p*_*s*_ (*t*) is the probability that the mutant lineage has not gone extinct, and *f*_*s*_ (*t*) is the average lineage size (conditioned on having not gone extinct):

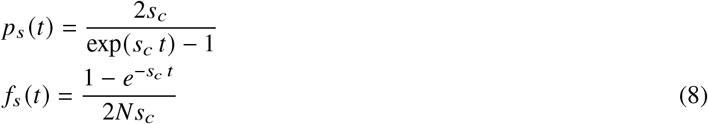

**So**, we can plug Eqn. 7 into Eqn. 6 to get:

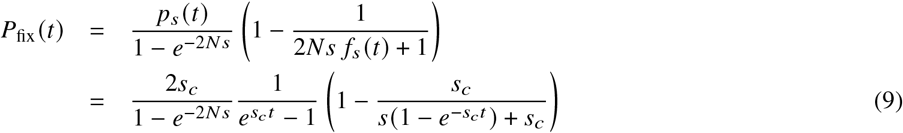

Proceed by sticking the above into Eqn. 5:

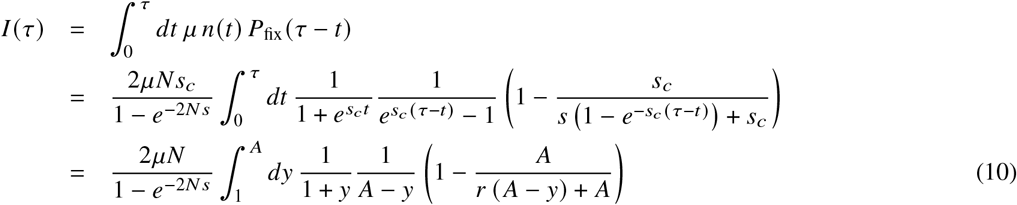

Here, 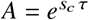 and *r* = *s*/*s*_*c*_, and we have changed variables to 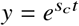. Evaluating the integral,

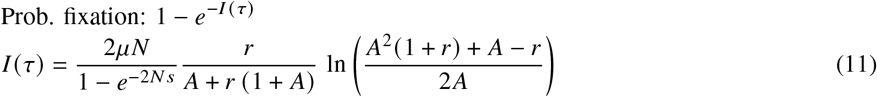

as reported in the main text.

### Numerical methods

All simulations are done with a simple implementation of Wright-Fisher dynamics with mutations and selection. We work with a fixed population *N* and multiple genotypes with *n*_*i*_ individuals each. At each time step, we draw the composition of the new population by sampling with replacement from the old one. The probability of sampling genotype *i* is *P*_*i*_ ∝ Σ _*j*_ *W*_*i j*_ *n* _*j*_, where:

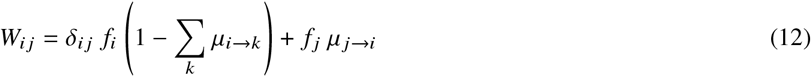

where *μ*_*i*→ *j*_ is the mutation rate from genotype *i* to genotype *j*.

These dynamics are implemented with the function *wrightFisherDynamics*, seen below. For instance, the code here simulates (many replicates of) a single mutator competition:

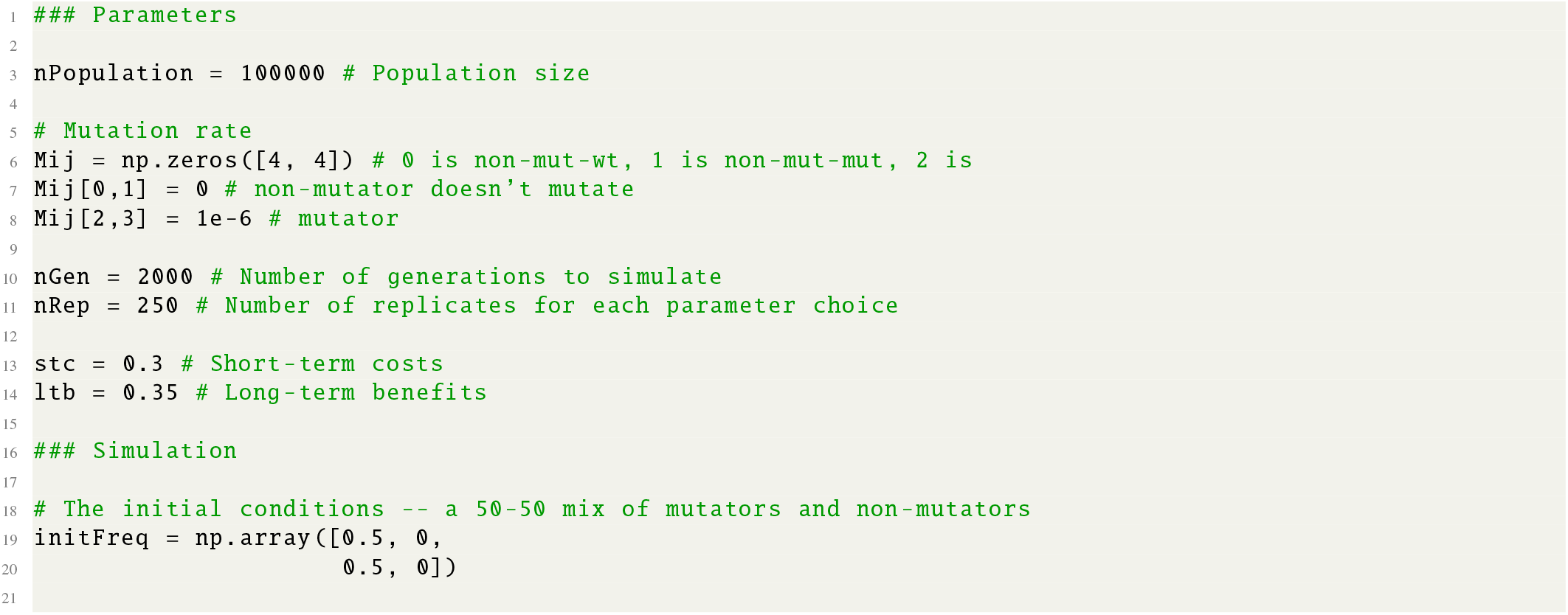
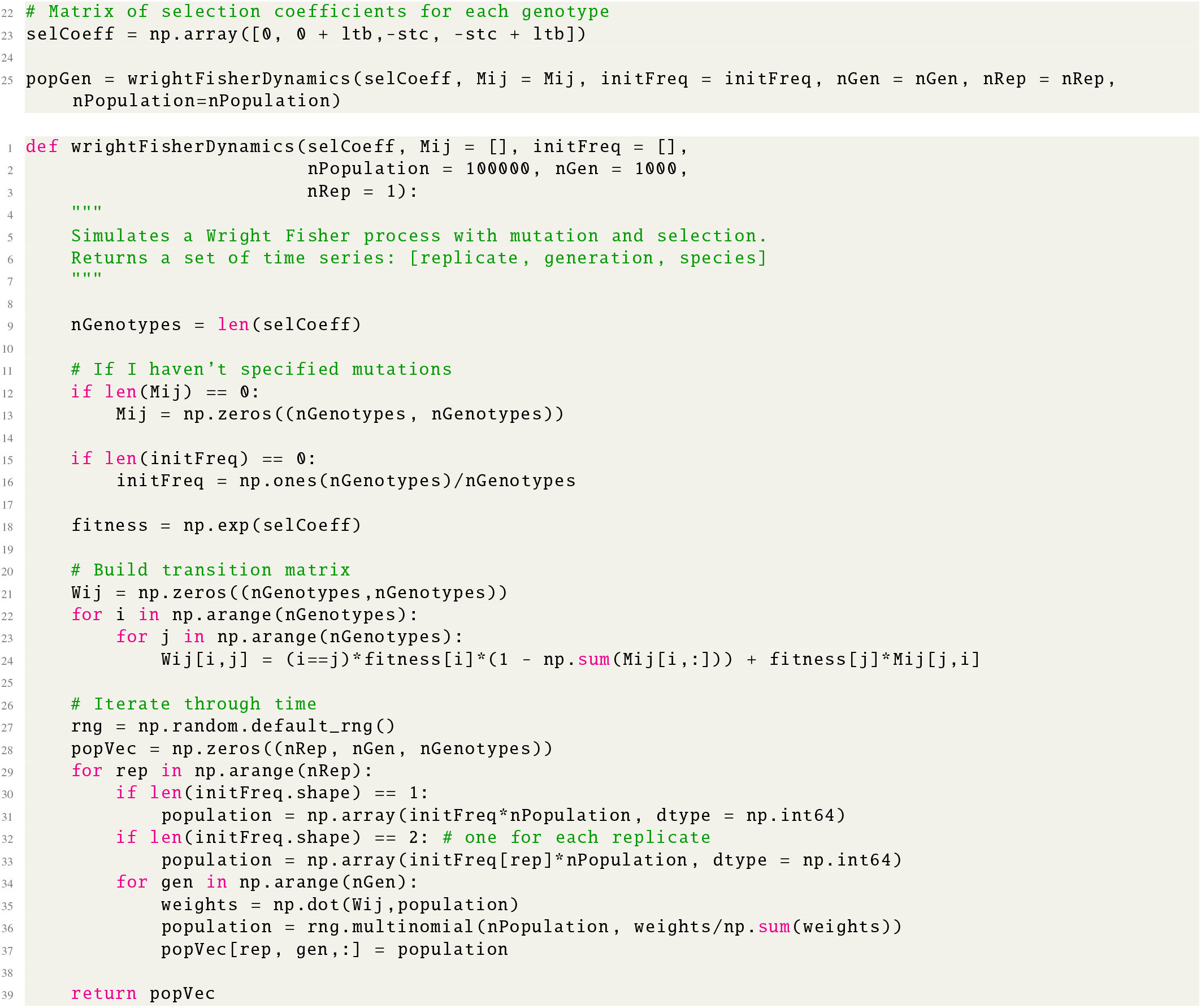

### Parameter values

- Fig. 1c of main text: population size *N* = 10^7^, non-mutator *μ* = 0, mutator *μ* = 10^−5^, strength of beneficial *s*_*b*_ ∈ {0.1, 0.15, 0.2, 0.25, 0.4}.
- Fig. 2e of main text: population size *N* = 10^5^, non-mutator *μ* = 0, mutator *μ* = 10^−5^. From left-to-right, pairs of {*s*_*c*_, *s*_*b*_} for the mutator are {10^−2^, 5 × 10^−3^ }, {10^−2^, 0.105}, {10^−2^, 0.015}, and {2 × 10^−3^, 10^−1^ }.
- Fig. 2f of main text: population size *N* = 10^5^, non-mutator *μ* = 0, mutator *μ* = 10^−6^. Data points were generated from a grid of values spanning *s*_*c*_ ∈ [0.05, 2] and *s*_*b*_ ∈ [0.01, 0.25], with 250 replicates for each pair of parameter values. Simulations were run for 2000 generations each.
- ‘Rock-paper-scissors’: Fig 5b of main text. We simulated 10,000 replicate Wright-Fisher simulations, 250 generations each, with the following parameters:
  1. A fixed long-term-benefit *s*_*b*_ = 0.3 for all strains.
  2. Non-mutator A: *μ* = 0, short-term cost *s*_*c*_ = 0.
  3. Mutator B: *μ* = 10^−5^, *s*_*c*_ = 0.08
  4. Super-mutator C: *μ* = 10^−3^, *s*_*c*_ = 0.2
- Time varying selection, Fig. 5d,e of main text: populations size *N* = 10^5^, non-mutator *μ* = 0, mutator *μ* = 10^−5^. All simulations began in an environment with *s*_*c*_ = 0.005 and *s*_*b*_ = 0. After time *τ*, populations were transitioned to an environment with an {*s*_*c*_, *s*_*b*_} of {0.5, 0.5025} (orange), {0.5, 0.51} (black), or {0.5, 0.55} (green).

## References

[1] Losos, J. B. The Princeton guide to evolution (Princeton University Press, 2017).

[2] Cairns, J., Overbaugh, J. & Miller, S. The origin of mutants. Nature 335, 142–145 (1988).

[3] Lynch, M. et al. Genetic drift, selection and the evolution of the mutation rate. Nat. Rev. Genet. 17, 704–714 (2016).

[4] Matic, I. et al. Highly variable mutation rates in commensal and pathogenic escherichia coli. Science 277, 1833– 1834 (1997).

[5] LeClerc, J. E., Li, B., Payne, W. L. & Cebula, T. A. High mutation frequencies among escherichia coli and salmonella pathogens. Science 274, 1208–1211 (1996).

[6] Loeb, L. A. Human cancers express mutator phenotypes: origin, consequences and targeting. Nature Reviews Cancer 11, 450–457 (2011).

[7] Sniegowski, P. D., Gerrish, P. J. & Lenski, R. E. Evolution of high mutation rates in experimental populations of e. coli. Nature 387, 703–705 (1997).

[8] Sane, M., Diwan, G. D., Bhat, B. A., Wahl, L. M. & Agashe, D. Shifts in mutation spectra enhance access to beneficial mutations. Proceedings of the National Academy of Sciences 120, e2207355120 (2023).

[9] Couce, A., Guelfo, J. R. & Blázquez, J. Mutational spectrum drives the rise of mutator bacteria. PLoS genetics 9, e1003167 (2013).

[10] Keightley, P. D. & Otto, S. P. Interference among deleterious mutations favours sex and recombination in finite populations. Nature 443, 89–92 (2006).

[11] Rutherford, S. L. & Lindquist, S. Hsp90 as a capacitor for morphological evolution. Nature 396 (1998).

[12] Johnson, M. S. & Desai, M. M. Mutational robustness changes during long-term adaptation in laboratory budding yeast populations. eLife 11, e76491 (2022).

[13] Woods, R. J. et al. Second-order selection for evolvability in a large escherichia coli population. Science 331, 1433– 1436 (2011).

[14] Kunkel, T. A. & Bebenek, K. DNA replication fidelity 1. Annu. Rev. Biochem. 69, 497–529 (2000).

[15] Barrick, J. E. & Lenski, R. E. Genome dynamics during experimental evolution. Nat. Rev. Genet. 14, 827–839 (2013).

[16] Drake, J. W., Charlesworth, B., Charlesworth, D. & Crow, J. F. Rates of spontaneous mutation. Genetics 148, 1667–1686 (1998).

[17] Zhu, Y. O., Siegal, M. L., Hall, D. W. & Petrov, D. A. Precise estimates of mutation rate and spectrum in yeast. Proceedings of the National Academy of Sciences 111, E2310–E2318 (2014).

[18] Jiang, P. et al. A modified fluctuation assay reveals a natural mutator phenotype that drives mutation spectrum variation within saccharomyces cerevisiae. Elife 10, e68285 (2021).

[19] Turajlic, S., Sottoriva, A., Graham, T. & Swanton, C. Resolving genetic heterogeneity in cancer. Nature Re-views Genetics 20, 404–416 (2019).

[20] Coelho, M. C., Pinto, R. M. & Murray, A. W. Heterozygous mutations cause genetic instability in a yeast model of cancer evolution. Nature 566, 275–278 (2019).

[21] Shor, E., Fox, C. A. & Broach, J. R. The Yeast Environmental Stress Response Regulates Mutagenesis Induced by Proteotoxic Stress. PLOS Genetics 9, e1003680 (2013).

[22] Habig, M., Lorrain, C., Feurtey, A., Komluski, J. & Stukenbrock, E. H. Epigenetic modifications affect the rate of spontaneous mutations in a pathogenic fungus. Nature Communications 12, 5869 (2021).

[23] Kimura, M. On the evolutionary adjustment of spontaneous mutation rates. Genetics Research 9, 23–34 (1967).

[24] Liberman, U. & Feldman, M. W. Modifiers of mutation rate: a general reduction principle. Theoretical population biology 30, 125–142 (1986).

[25] Dawson, K. J. Evolutionarily stable mutation rates. Journal of theoretical biology 194, 143–157 (1998).

[26] Lynch, M. The cellular, developmental and population-genetic determinants of mutation-rate evolution. Genetics 180, 933–943 (2008).

[27] Desai, M. M. & Fisher, D. S. The balance between mutators and nonmutators in asexual populations. Genetics 188, 997–1014 (2011).

[28] Lynch, M. The lower bound to the evolution of mutation rates. Genome biology and evolution 3, 1107–1118 (2011).

[29] Soderberg, R. J. & Berg, O. G. Kick-starting the ratchet: The fate of mutators in an asexual population. Genetics 187, 1129–1137 (2011).

[30] James, A. & Jain, K. Fixation probability of rare non-mutator and evolution of mutation rates. Ecology and Evolution 6, 755–764 (2016).

[31] Good, B. H. & Desai, M. M. Evolution of mutation rates in rapidly adapting asexual populations. Genetics 204, 1249–1266 (2016).

[32] Ferrare, J. T. & Good, B. H. Evolution of evolvability in rapidly adapting populations. Nature Ecology & Evolution (2024).

[33] Gunge, N. & Sakaguchi, K. Intergeneric transfer of de-oxyribonucleic acid killer plasmids, pgkl1 and pgkl2, from kluyveromyces lactis into saccharomyces cerevisiae by cell fusion. Journal of bacteriology 147, 155–160 (1981).

[34] Ravikumar, A., Arrieta, A. & Liu, C. C. An orthogonal DNA replication system in yeast. Nat. Chem. Biol. 10, 175–177 (2014).

[35] Ravikumar, A., Arzumanyan, G. A., Obadi, M. K. A., Javanpour, A. A. & Liu, C. C. Scalable, continuous evolution of genes at mutation rates above genomic error thresholds. Cell 175, 1946–1957.e13 (2018).

[36] Ravasio, R. et al. A minimal scenario for the origin of non-equilibrium order (2024). URL http://arxiv.org/abs/2405.10911.

[37] Payne, J. L. & Wagner, A. The causes of evolvability and their evolution. Nature Reviews Genetics 20, 24–38 (2019).

[38] Barnett, M., Zeller, L. & Rainey, P. B. Experimental evolution of evolvability. bioRxiv 2024–05 (2024).

[39] Desai, M. M., Fisher, D. S. & Murray, A. W. The speed of evolution and maintenance of variation in asexual populations. Current biology 17, 385–394 (2007).

[40] McDonald, M. J., Hsieh, Y.-Y., Yu, Y.-H., Chang, S.-L. & Leu, J.-Y. The evolution of low mutation rates in experimental mutator populations of saccharomyces cerevisiae. Current Biology 22, 1235–1240 (2012).

[41] Wei, W. et al. Rapid evolution of mutation rate and spectrum in response to environmental and population-genetic challenges. Nature communications 13, 4752 (2022).

[42] Stark, M. & Boyd, A. The killer toxin of Kluyveromyces lactis: characterization of the toxin subunits and identification of the genes which encode them. The EMBO Journal 5, 1995–2002 (1986).

[43] Klassen, R. & Meinhardt, F. Linear protein-primed replicating plasmids in eukaryotic microbes. In Microbial Lin-ear Plasmids, 187–226 (Springer, 2007).

[44] Hopfield, J. J. Kinetic proofreading: a new mechanism for reducing errors in biosynthetic processes requiring high specificity. Proceedings of the National Academy of Sciences 71, 4135–4139 (1974).

[45] Ninio, J. Kinetic amplification of enzyme discrimination. Biochimie 57, 587–595 (1975).

[46] Phillips, R. The molecular switch: Signaling and Allostery (Princeton University Press, 2020).

[47] Rix, G. et al. Continuous evolution of user-defined genes at 1-million-times the genomic mutation rate (2023). URL https://www.biorxiv.org/content/10.1101/2023.11.13.566922v1.

[48] Lozovsky, E. R. et al. Stepwise acquisition of pyrimethamine resistance in the malaria parasite. Proceedings of the National Academy of Sciences 106, 12025–12030 (2009).

[49] Sirawaraporn, W., Sathitkul, T., Sirawaraporn, R., Yuthavong, Y. & Santi, D. V. Antifolate-resistant mutants of plasmodium falciparum dihydrofolate reductase. Proceedings of the National Academy of Sciences 94, 1124–1129 (1997).

[50] Fields, S. The two-hybrid system to detect protein-protein interactions. Methods 5, 116–124 (1993).

[51] Bennett, C. H. Dissipation-error tradeoff in proofreading. BioSystems 11, 85–91 (1979).

[52] Banerjee, K., Kolomeisky, A. B. & Igoshin, O. A. Eluci-dating interplay of speed and accuracy in biological error correction. Proceedings of the National Academy of Sciences 114, 5183–5188 (2017).

[53] Mallory, J. D., Kolomeisky, A. B. & Igoshin, O. A. Trade-offs between error, speed, noise, and energy dissipation in biological processes with proofreading. The Journal of Physical Chemistry B 123, 4718–4725 (2019).

[54] Midha, T., Mallory, J. D., Kolomeisky, A. B. & Igoshin, O. A. Synergy among pausing, intrinsic proofreading, and accessory proteins results in optimal transcription speed and tolerable accuracy. The Journal of Physical Chemistry Letters 14, 3422–3429 (2023).

[55] Murugan, A., Huse, D. A. & Leibler, S. Speed, dissipation, and error in kinetic proofreading. Proceedings of the National Academy of Sciences 109, 12034–12039 (2012).

[56] Raynes, Y., Wylie, C. S., Sniegowski, P. D. & Weinreich, D. M. Sign of selection on mutation rate modifiers depends on population size. Proceedings of the National Academy of Sciences 115, 3422–3427 (2018).

